# Evaluating single-subject study methods for personal transcriptomic interpretations to advance precision medicine

**DOI:** 10.1101/428581

**Authors:** Samir Rachid Zaim, Colleen Kenost, Joanne Berghout, Helen Hao Zhang, Yves A. Lussier

## Abstract

**Background:** Gene expression profiling has benefited medicine by providing clinically relevant insights at the molecular candidate and systems levels. However, to adopt a more ‘precision’ approach that integrates individual variability including ‘omics data into risk assessments, diagnoses, and therapeutic decision making, whole transcriptome expression analysis requires methodological advancements. One need is for users to confidently be able to make individual-level inferences from whole transcriptome data. We propose that biological replicates in isogenic conditions can provide a framework for testing differentially expressed genes (**DEGs**) in a single subject (**ss**) in absence of an appropriate external reference standard or replicates.

**Methods:** Eight ss methods for identifying genes with differential expression (**NOISeq, DEGseq, edgeR, mixture model, DESeq, DESeq2, iDEG, and ensemble**) were compared in **Yeast** (parental line versus snf2 deletion mutant; n=42/condition) and **MCF7** breast-cancer cell (baseline and stimulated with estradiol; n=7/condition) RNA-Seq datasets where replicate analysis was used to build reference standards from NOISeq, DEGseq, edgeR, DESeq, DESeq2. Each dataset was randomly partitioned so that approximately two-thirds of the paired samples were used to construct reference standards and the remainder were treated separately as single-subject sample pairs and DEGs were assayed using ss methods. Receiver-operator characteristic (**ROC**) and precision-recall plots were determined for all ss methods against each RSs in both datasets (525 combinations).

**Results:** Consistent with prior analyses of these data, **~**50% and ~15% DEGs were respectively obtained in Yeast and MCF7 reference standard datasets regardless of the analytical method. NOISeq, edgeR and DESeq were the most concordant and robust methods for creating a reference standard. Single-subject versions of NOISeq, DEGseq, and an ensemble learner achieved the best median ROC-area-under-the-curve to compare two transcriptomes without replicates regardless of the type of reference standard (>90% in Yeast, >0.75 in MCF7).

**Conclusion:** Better and more consistent accuracies are obtained by an ensemble method applied to single-subject studies across different conditions. In addition, distinct specific sing-subject methods perform better according to different proportions of DEGs. Single-subject methods for identifying DEGs from paired samples need improvement, as no method performs with both precision>90% and recall>90%. http://www.lussiergroup.org/publications/EnsembleBiomarker

## Background

Gene expression profiling has benefited medicine by characterizing cellular states throughout development and differentiation, describing the pathological processes occurring during disease, and providing clinically relevant insights at the molecular candidate and systems levels. As medicine moves to adopt a more ‘precision’ approach that integrates individual variability including ‘omics data into risk assessments, diagnoses, and therapeutic decision making, whole transcriptome expression analyses using technologies such as RNA-Seq are poised to become foundational methods(1). Still, there are issues to resolve before this promise can be realized; most related to data analysis and interpretation rather than data collection, though all areas can still be better optimized. Major areas for computational analytical methods improvements include (i) the development of a well-validated reference standard, thoroughly vetted and solidly benchmarked for a given investigation, and (ii) the ability to confidently make individual-level inferences from transcriptomic data.

To the last point, the majority of *differentially expressed gene* (**DEG**) analysis methods currently available have been designed to make inferences at the population level about diseases or conditions, not for individual patients. These experiments and analytical approaches seek to define and characterize the common and consensus processes that differentiate or underlie two (or more) states. In basic research using model organisms, researcher control over genotype and experimental parameters allow genotype-level inference by using a two-group comparison that leverages three or more replicates per group(2). In clinical research using human subjects, however, the genotypic and lived experience diversity of each subject introduces substantial biological variability and noise into expression data. This then requires tens to thousands of genotype-distinct replicate samples to draw inferences about the population(s) and condition(s) of interest, but simultaneously ignores or prohibits individual-level variation and inferences unless they can be classified according to stratification patterns common enough to be noticed(3). To adapt the tools designed for populations into tools appropriate for individual-level inference requires either the use of replicates (mimicking the style of a model organism experiment and reducing the cross-sample noise to primarily stochastic and technical factors), *a priori* distribution and parameter assumptions, or data-derived models to create an expected distribution useful for comparison. However, in practice, it is not cost-effective and often entirely infeasible to obtain replicate samples from the clinic procedures. Since DEG analysis methods were validated using replicates(3, 4), there remains a need to learn how well a DEG method designed for identifying differential expression would perform in real-world conditions and when replicates are unavailable (**ss-DEG Methods**).

Novel methodological advances designed with single subjects in mind have begun to be proposed(3, 4). While accurately discovering DEGs between two RNA-Seq samples remains a challenge and insufficiently studied(3, 4), methods identifying *differentially expressed gene sets and pathways* between two transcriptomes applicable to single-subject studies have been reproducibly demonstrated as feasible(3, 4) in simulations(5), retrospective studies in distinct datasets (5–10), cellular assays (11, 12), as well as in one clinical classifier (13) (**details in supplement Table S1**). These comprehensive validations of gene set/pathway-level methods established the feasibility of single-subject interpretation of the transcriptomes and stimulate further investigations to improve more precise methods for determining the underlying differentially expressed genes. However, transcriptional dynamics operating and validated at the gene set or pathway-level cannot straightforwardly be deconvoluted to identify specific transcripts altered in a single-subject. A recent study provides a comparison of accuracy of five ss-DEGs methods using computer simulations of several data models with genomic dysregulation ranging from 5 to 40% DEGs(14). A partial biological validation was conducted for one ss-DEG methods, NOIseq(15), confirming the top 400 DEG signals by qPCRs. Yet, and to the best of our knowledge, no study has comprehensively validated nor compared the accuracies of ss-DEG methods using biological or clinical datasets on a transcriptome scale. In addition, no framework has been proposed on how to conduct such a comprehensive validation.

We and others(16) propose that there is a knowledge gap in the field with regards to optimizing the operating characteristics of the state-of-the-art RNA-Seq analytics for precision medicine: what are the best ss-DEG methods for interrogating two RNA-Seq samples from one patient taken in two different conditions without replicates? Reliable and accurate of ss-DEG methods can have practical utility. For example, the comparison of affected versus unaffected samples (e.g., cancer v. non-cancer) can provide valuable insight into the genetic variables involved in a disease’s pathophysiology and therapeutics. Similarly, using a patient’s healthy tissue as its baseline to compare treated tissue or evolution over time provides another framework to design analytics and assays for precision medicine.

We thus designed this study under the following premise: *isogenic biological replicates (genome matched) conditions can provide a framework for testing single-subject methods in absence of an external reference standard*. In this study, we aim to identify the best-performing techniques and parameters in absence of replicates of distinct single-subject (**ss**) methods predicting differentially expressed genes (**DEGs**). In addition, we hypothesized, implemented, and evaluated an ensemble method as possibly more robust across different conditions of application for determining DEGs in single subjects.

## Methods

Figure 1 provides an overview of the experimental design, including the methods and recommendation for using an ensemble learner approach to develop robust reference standards in ss studies.

**Figure 1.**
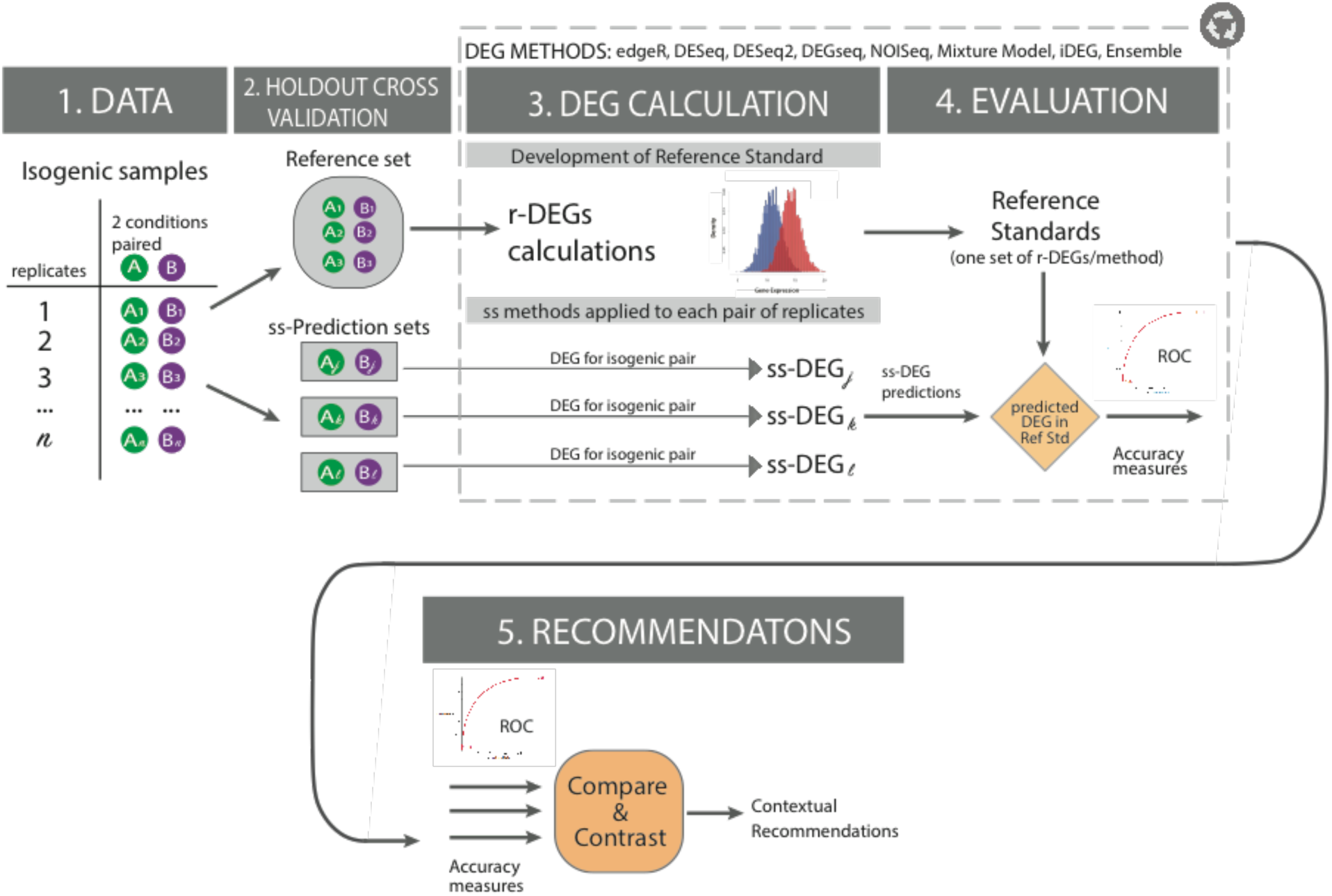
Overview of the evaluation strategy of methods designed for transcriptome analysis in a single subject from paired samples. *Motivation:* Identifying the gene products altered between two conditions in a single subject without replicates **(ss-DEGs)** is highly relevant in precision medicine and is the underlying motivation of this study. While conventional analytical methods, designed to find coordinated averages between two groups of subjects, may be applied to discover differences between isogenic *replicates* studied in distinct conditions (**r-DEGs**), precision medicine has helped usher in the possibility that diagnosis, prognosis, and therapeutic choices may come to be determined *more accurately from single-subject measurements*. For example, accurate ss-DEGs methods could be applied to study (i) cancer vs unaffected adjacent tissue or (ii) an *ex-vivo* cellular provocation assay operating on relevant tissue with or without therapy. ***Description of the evaluation framework*. Step 1 DATA**. A dataset comprising multiple biological replicates of isogenic transcriptomes observed on samples taken in distinct biological conditions is identified. **Step 2 HOLDOUT CROSS-VALIDATION**. The replicates are split into two groups of independent samples: a reference set and a single-subject (**ss**) prediction set. **Step 3 DEG CALCULATIONS**. Each r-DEG method (e.g., EdgeR, DESeq, etc.) is applied independently to the reference set to generate multiple reference standards, as each method has biases and none can be truly considered as a gold standard (**Step 3, top panel**). The reference set consists of biological replicates between two conditions of isogenic samples, and is thus a proxy for studying and mimicking the biologic variation of one subject (e.g., isogenic) and each set of r-DEGs is an attempt at becoming a gold standard. In parallel, each ss-DEG method is applied to independent pairs of samples (one in each condition) taken from the prediction set, each as a proxy to a single subject (**Step 3. Bottom panel**). **Step 4 EVALUATION**. Accuracy scores are determined for each ss-DEG method against each r-DEG-derived reference standard. To avoid biases, an ss-DEG method applied on paired sample of the prediction set is never evaluated against its corollary application as well as a r-DEG method against the reference set. **Step 5 RECOMMENDATIONS**. Summary statistics are conducted across all experiments to determine the best ss-DEGs according to the conditions of application.

### Computing Environment

All analyses in this study were conducted in the R programming language, using R 3.4.0(17), and all the code is freely available at http://www.lussiergroup.org/publications/EnsembleBiomarker.

### Datasets

In this study, two distinct isogenic RNA-Seq datasets were used to calculate the reference standards and to conduct the single-subject studies. Both datasets have previously been validated as reference standards to evaluate methods that determine differentially expressed genes (**DEGs**) from RNA-Seq, using cohort or groups of biological replicates (**r-DEGs methods**) rather than for determining the accuracy of single-subject DEGs (**ss-DEGs methods**) of the current study.

**<u>Yeast dataset:</u>** The first dataset (hereinafter) “Yeast” is comprised of 48 wild-type (BY4741 strain, WT) or **Δ**snf2 mutant biological yeast replicates *(Saccharomyces cerevisiae)* generated on the same background. RNA-Seq analysis and mapping includes 7,126 measured genes (18). We followed the author’s data preprocessing guidelines and conducted our studies using their suggested 42 WT and 44 **Δ**snf2 ‘clean’ replicates. Normalized and preprocessed data were downloaded as prepared by the original authors from their GitHub^i^ repository, under their “Preprocessed_data” directory. 48 expression count files were downloaded for the two conditions, respectively, retaining the “clean” replicates for analysis. <u>**MCF7** dataset</u>: Our second dataset consists of 7 biological replicates of human MCF7 cells (~ 22,000 measured genes) which were either treated with 10 nM 17β-estradiol (E2) or cultured as unstimulated controls (19). We used the 30M read replicates available in the MCF7 dataset, which is available open source online under the Gene Expression Omnibus repository (20) (id = GSE51403). Normalized and preprocessed were downloaded on January 21, 2018.

### Preprocessing and Prediction Set Construction

The Yeast and MCF7 datasets for the biological validation studies were used entirely as obtained, with no additional pre-processing steps or data manipulation. Transcript mapping, filtering, normalization, and batch correction details can be found in the original publications (18, 19). In the MCF7 dataset, the following 4 biological replicates (“565–576”,”564–572”,”566–570”,”562–574”) were randomly selected and set aside as the training set, and the additional 3 (“563–577”,”568–575”,”569–571”) were then used to construct and evaluate how well the DEG methods could recapture the signal. Similarly, in the Yeast dataset, 30 replicates were randomly selected and set aside to construct the reference standard, and the remaining available 12 ‘clean’ replicates were used in the single-subject studies.

### DEG Methods

The study is designed to better understand how single-subject studies can be conducted in biological and clinical precision medicine settings, where a true gold standard accurately reflective of a known ground truth does not always exist. To this end, we compared published and novel computational methods designed to detect DEGs from <u>single-subject without replicates (**ss**)</u> with a variety of well-validated and widely-used RNA-Seq analysis methods designed to identify DEGs from <u>cohort or replicate-based comparisons (**r-DEG**)</u> (**Table 1**)(5, 15, 21–25). With the exception of NOISeq that has been directly designed for application to a single-subject under two conditions without replicates (NOISeq-sim implementation), the other replicate-based methods (**Table 1**) have not been designed nor systematically tested for accurate performance in single-subject paired-sample condition where replicates are not available. However, for the methods we have selected, the authors have estimated the required parameters to perform these comparisons, which are included in package documentation. All methods were implemented according to the default parameters provided for isogenic conditions (genotype-replicates) in the original publications. For NOISeq, we used noiseqbio under their default parameter settings to generate the reference standard, and noiseq-sim (setting the parameters replicates = “no” and nss=3) for the single-subject studies. For DESeq, in the estimateDispersions function, the method parameter is set to ‘per-condition’ for the replicated study, and ‘blind’ for the single-subject studies. For edgeR, we use the “genetically identical model organisms” replicate-type in order to set the appropriate BCV value; and finally, DEGseq and DESeq2 are implemented in wrapper functions using their default parameters.

### ss-DEG calculations

For each DEG method in **Table 1**, we calculated ss-DEGs for 12 distinct pairs of samples in the Yeast datasets, and 3 pairs of samples in the MCF7 dataset by sampling without replacement to create independent pairs. Of note, while many of the methods were not intended nor validated for ss-DEG calculations, the authors of each of the r-DEG methods listed in Table 1 did indicate their possible application to two-sample comparisons and provided unpublished approaches to adapt or estimate the parameters required for such processing. All details and code are available at http://www.lussiergroup.org/publications/EnsembleBiomarker.

False Discovery Rates (FDRs) were calculated using Benjamini-Yekutieli (26). Mixture Models were implemented as described by Li et al (5) and a posterior probability rather than a FDR is utilized for the receiver-operator characteristics curves and the precision-recall plots. In **Figures 4 and 5**, the posterior probability >95% of a fold change between two samples being a significant DEG was utilized as a Mixture Model cutoff corresponding to the FDR<5%.

**Table 1.**
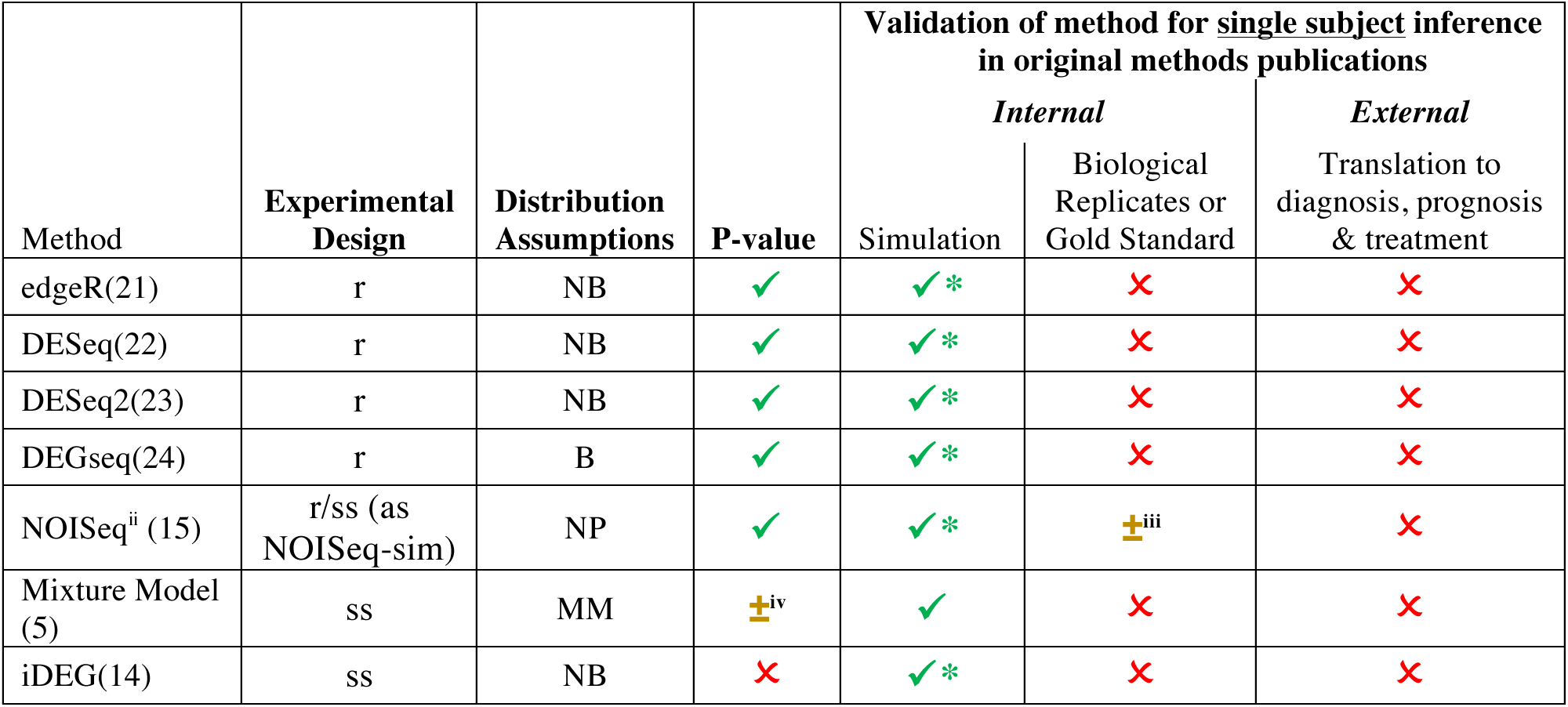
DEG Methods for Single-Subject Studies and their previous validations.^ii^^iii^^iv^. Legend: NB= Negative Binomial, B= Binomial, NP= Non-Parametric, MM= Mixture Model, ss= single-subject analytics, r= analytics of between group of replicates, ✓ = completed, ± = partially addressed, ✘ = not addressed

### Developing an Ensemble Learner across ss-DEG methods

Since differences across individual techniques showed variable performance, we constructed a naïve **ensemble** predictor (hereinafter referred to as the “ensemble”). Continuing to treat each independent single-subject as an independent assay, the ensemble combined ss-DEG predictions from DEGSeq, NOISeq, mixture models, and edgeR by taking the arithmetic mean of the FDR corrected values. Since the single-subject implementations of both DESeq and DESeq2 had extremely low recall (recall<1% of DEGs; Results, Figures 3–5), these were excluded from the ensemble. Finally, since iDEG(14) is a preprint publication, we decided against including them it in the ensemble in order to create an ensemble consisting exclusively of published and peer-reviewed techniques. We adopted the strategy used by random forests(27) in constructing an ensemble (e.g., average the predictions of individual decision trees into an ensemble predictor), since random forests have shown a great level of success in genomics.

Therefore, the ensemble was constructed using the following formula:

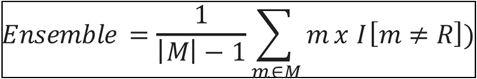

where M is the set of models used to build the ensemble (i.e., M = {DEGseq, NOISeq, mixture models, edgeR}), |M| is the cardinality of M (e.g., the number of models), *I* is the indicator function, and R is the method used to build the reference standard. For example, when edgeR was used to build the replicated reference standard, the ensemble omits edgeR as follows.

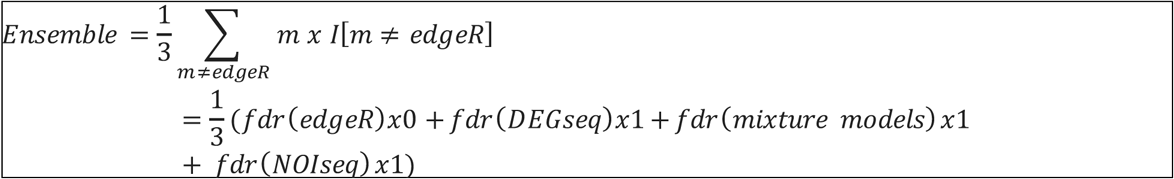

The reference standard is omitted from the construction of the ensemble in order to minimize any of its potential biases or unfair advantages.

### Reference Standard Construction

Each r-DEGs method in **Table 1** was used to construct a reference DEG standard once using n=30 wild type versus n=30 snf2 mutant yeast for the Yeast dataset, and n=4 unstimulated vs n=4 estrogen-stimulated in the MCF7 dataset. DEGs identified by each r-DEG method were compared against one another to assess cross-method overlap to quantify the variability and reliability of reference standards. All r-DEG methods were implemented using their recommended default settings as described earlier in the “DEG Methods” section.

In the original manuscript describing the MCF7 dataset (19), the authors set a threshold resulting in approximately 3,300 genes detected as DEGs by edgeR when all 7 replicates were used. Therefore, we adjusted our False Discovery Rate (**FDR**)(28) thresholds in each method to operate similarly and detect approximately 3,300 DEGs (~ 15% of genes). In the Yeast dataset, we mimicked the authors’ experimental design and set our FDR-thresholds for DEG detection at FDR<5%, which resulted in a varying number of DEGs per method that closely resembled the results (e.g., number of DEG calls) obtained by the original authors analysis of this dataset. **Table 2** summarizes the operating characteristics of these methods in both datasets.

### Evaluation

Each of the single-subject methods in **Table 1** were evaluated in single-subject studies using the nonoverlapping remainder of replicates as a prediction set (twelve pairs of single-subject samples for Yeast and three for MCF7) using Precision-Recall (**PR**) and receiver-operator characteristic (**ROC**) plots. In **Figure 3**, when a method from **Table 1** was used to construct a reference standard, the single-subject implementation of that same method was omitted from that series of analyses (i.e., in the reported summary statistics of accuracies of ss-DEGseq, the reference standard based on DEGseq was omitted from all precision-recall and accuracy metric evaluations).

Our prediction set of three single-subject studies in the MCF7 dataset and twelve single-subject studies in Yeast, together with five methods for replicate-derived reference standards, lead to fifteen (MCF7) + 60 (Yeast) sets of precision-recall (PR) and ROC curves (see **Figure 3** for an illustrative example). Therefore, in order to meaningfully evaluate the methods across all conditions, we summarized each technique’s performance by analyzing their area under the curve (**AUC**), by calculating the AUCs in the PR and ROC curves. Furthermore, we illustrate each method’s operating characteristics by creating Precision-Recall confidence regions which are 1-standard deviation bands around their mean precision and recall, at FDR = 5%, 10%, and 20% (1% also calculated, not shown).

In **Figures 4 and 5**, the Union and Intersection of reference standards **(Table 2)** were utilized to establish the summaries of accuracies. The PR and ROC plots were generated using the *precrec* R package (29) and the boxplots were created using the *ggplot2* (30) graphics library in R.

**Table 2.**
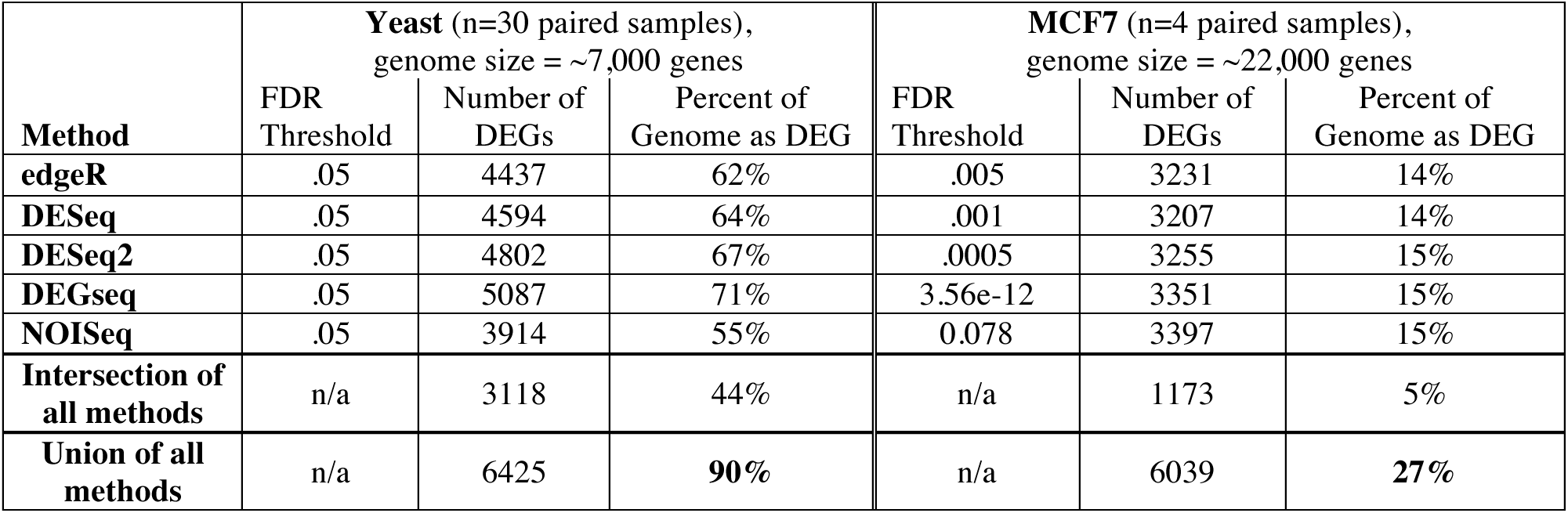
Operating Characteristics for each r-DEG methods applied to datasets with replicates to generate Reference Standards. FDRs are adjusted to obtain lists of DEGs of the same length as reported in the original publications. As shown with the intersection of all DEGs predicted by distinct methods, determining a gold standard in RNA-Seq analyses of multiple biological isogenic replicates remains a challenge. n/a = not applicable.

| Method | Yeast (n=30 paired samples),<br>genome size = ~7,000 genes |  |  | MCF7 (n=4 paired samples),<br>genome size = ~22,000 genes |  |  |
| --- | --- | --- | --- | --- | --- | --- |
|  | FDR<br>Threshold | Number of<br>DEGs | Percent of<br>Genome as DEG | FDR<br>Threshold | Number of<br>DEGs | Percent of<br>Genome as DEG |
| edgeR | .05 | 4437 | 62% | .005 | 3231 | 14% |
| DESeq | .05 | 4594 | 64% | .001 | 3207 | 14% |
| DESeq2 | .05 | 4802 | 67% | .0005 | 3255 | 15% |
| DEGseq | .05 | 5087 | 71% | 3.56e-12 | 3351 | 15% |
| NOISeq | .05 | 3914 | 55% | 0.078 | 3397 | 15% |
| Intersection of<br>all methods | n/a | 3118 | 44% | n/a | 1173 | 5% |
| Union of all<br>methods | n/a | 6425 | **90%** | n/a | 6039 | **27%** |

## Results

Evaluating DEGs between two conditions without replicate was not previous conducted using biologic samples. As previously reported by other authors, constructing a reliable reference standard from RNA-seq analytic methods remains a challenge (31) to date even in presence of 30 replicates in each condition as in our Yeast dataset. As shown in **Figure 2**, NOISeq, edgeR and DESeq were the most concordant and robust methods for creating a reference standard. However, the overall concordance between all methods varies substantially (**Table 2**). For example, the authors of the original Yeast dataset report ~60% DEGs, while the union of all methods can find as many as 90% DEGs, but their intersection report a mere 44%.

Since no single reference standard is truly a statement of truth, nor their union or intersection, we systematically evaluated methods discovering DEGs in two conditions without replicates against all reference standards. As discussed in the methods, distinct samples were utilized for calculating the reference standard and for estimating DEGs between paired transcriptomes. **Figure 3** demonstrates nine out of the possible 420 PR and ROC curve combinations for the Yeast dataset (5 reference standards x 12 independent sets of two paired samples × 7 methods evaluations in estimating DEGs from two conditions without replicates). The 420 Yeast PR and ROC plots and the 105 MCF7 PR and ROC plots are respectively summarized in **Panel A** of **Figures 3** and **4**. **Panel B** of **Figures 3** and **4** provide point predictions of each method at specific FDRs. The results are also illustrated in PR and ROC curves, one for each FDR cutoff (FDR = 5%, 10%, and 20%). The confidence regions in **Panels B** show how each of the methods performed across all samples, by providing their mean precision and recall, as well as a 1-SD band above/below and right/left of its mean precision-recall coordinate.

As FDR increases, the techniques increase their recall at the expense of some precision, with the exception, of DEGseq whose precision and recall in the Yeast dataset minimally increases. DEG detection methods like Mixture Model and DEGseq perform fairly consistently across all samples, resulting in narrower confidence regions whereas NOISeq and iDEG’s variability lies on the higher end of the spectrum. Note, DESeq2 is not shown in **Panel B** neither in **Figure 4** nor in **Figure 5** given its failure to produce any predictions at the selected FDR cutoffs.

**Figure 2:**
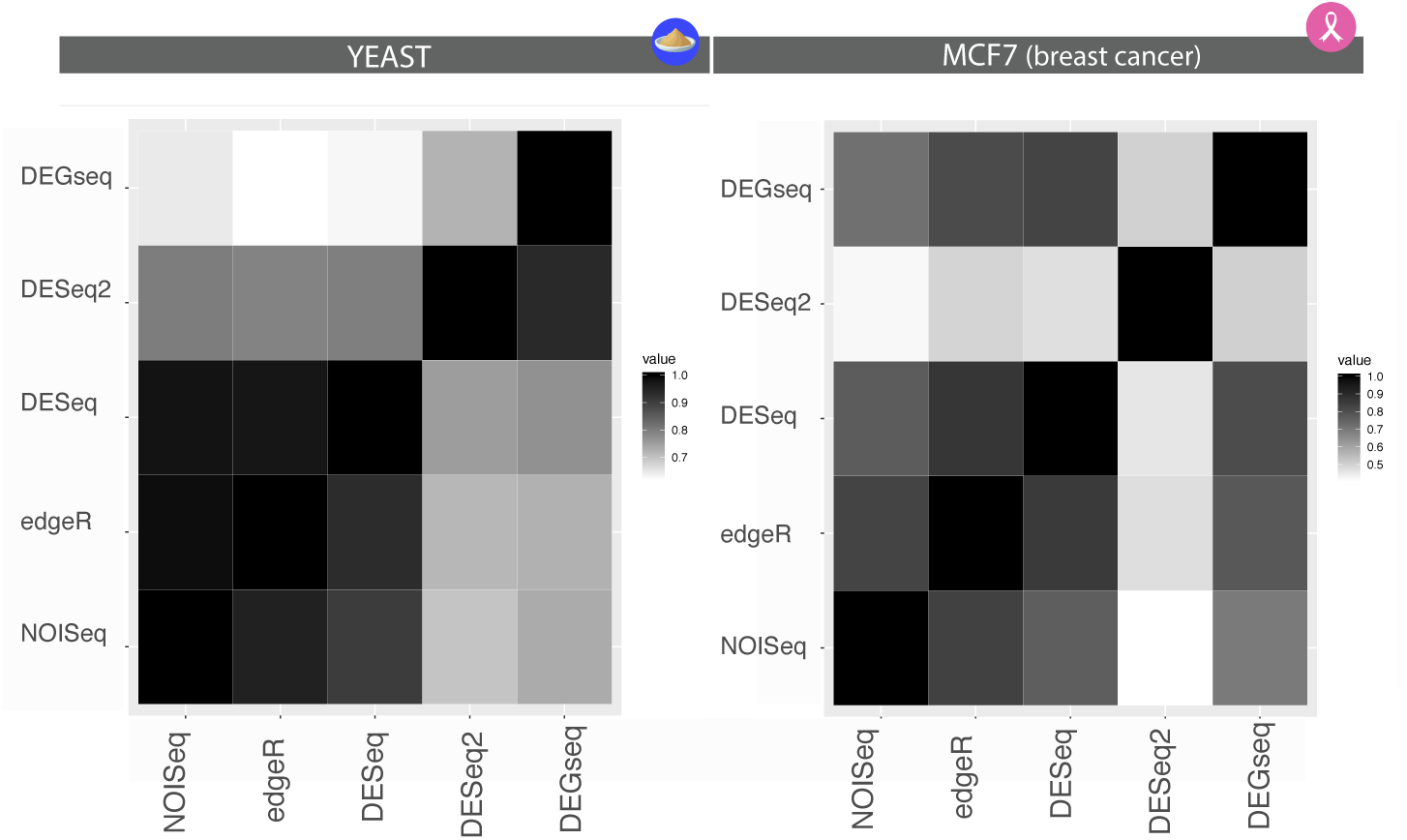
Comparison of Reference Standards demonstrates robust concordance between NOISeq, edgeR, and DESeq as well as inconsistent overlaps among other combinations. Each method’s pairwise concordance with one another (identity overlap of DEGs) is shown, with the diagonal entries as the total number of DEGs of each respective method, demonstrating the vulnerability of studies relying on a single method to develop a reference standard. The pairwise intersections were calculated using the count of DEGs in the methods of each column as the denominator. The heatmap is approximately, but not exactly, symmetric given the different denominators of comparing edgeR’s intersection with NOISeq vs. comparing NOISeq’s intersection with edgeR. In both Yeast (n=30) and MCF7 (n=4), edgeR, NOIseq, and DESeq show the best concordance to one another, while DESeq2 has the least concordance to any other method. DESeq2 shows the lack of agreement between what it considers DEGs and the rest of the methods, whereas in the left panel, both DESeq2 and DEGseq differentiate themselves from the cohort. This highlights the need for a consensus as some methods might make certain DEG calls that other methods miss and vice-versa. A conservative approach would be the intersection of all whereas an anti-conservative approach would take the union.

**Figure 3.**
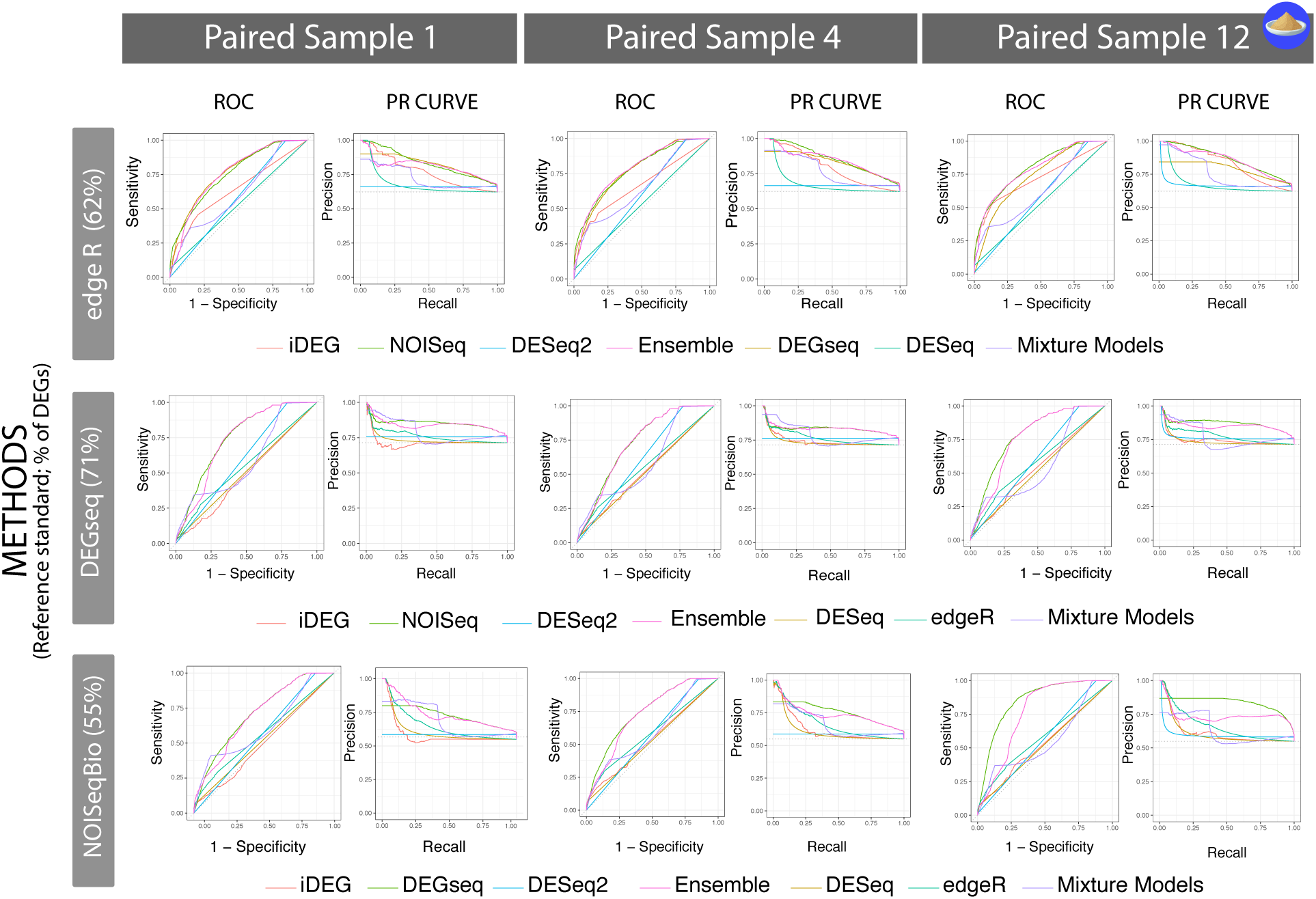
Exemplar accuracies of seven types of ss-DEGs methods validated against three rs-DEG-derived reference standards indicate a high-level of variability across ss-DEG methods and across reference standards, as well as a low to moderate-level of variability within ss-DEG methods and between biological replicates. The Precision-Recall and ROC curves across individual samples (Yeast) show that even in isogenic settings, a fair amount of biological variability exists. Furthermore, these single-subject studies provide a thorough comparison of each ss-DEG method’s performance and consistency in absence of replicates, allowing us to understand which tools have a greater potential for advancing precision medicine. For example, in the Yeast dataset, under >40 biological replicates, the authors recommended DESeq and DESeq2. However, in absence of biological replicates, these techniques performed conservatively and, on average, the worst.

**Figure 4.**
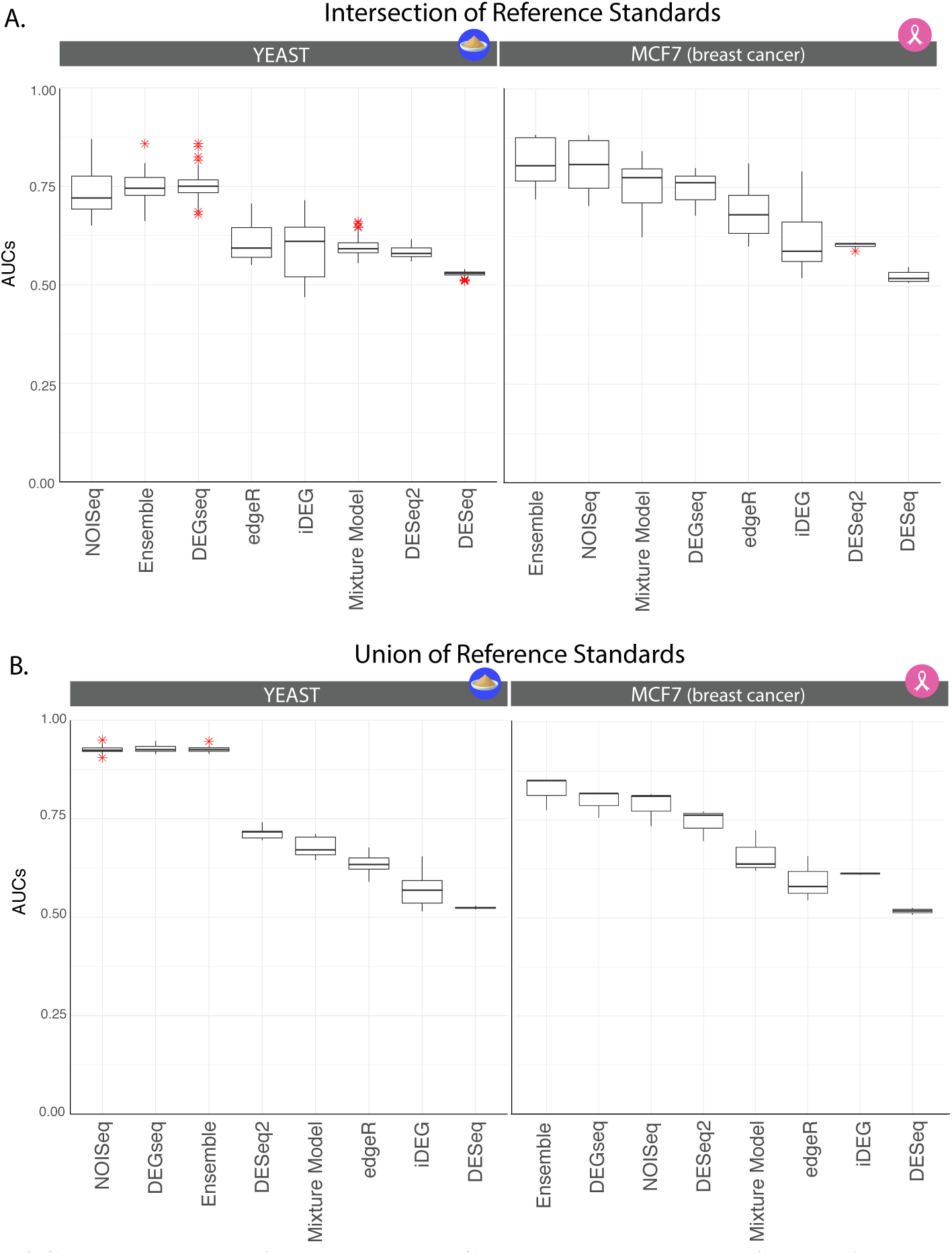
The ROC summary plots in Yeast and MCF7 show a spectrum of operating characteristics and performance across techniques, indicating the need for an ensemble-like approach. The Yeast case study produced reference standards that predicted between 55% and 70% of the genes in the genome as DEGs, while the MCF7 breast cancer cell lines predicted ~15% DEGs. In **Panel A**, the overtly-conservative scenario, the reference standard was constructed by taking <u>intersection</u> of the DEG lists from cohort analysis of the dataset with DESeq2, DEGSeq, edgeR, NOISeq-BIO (6425 genes as DE), whereas in the anti-conservative scenario (**Panel B**), the reference standard was constructed taking the <u>union</u> of all techniques (6039 genes as DE). The anti-conservative scenario facilitates the prediction task as a larger number of genes are called DEGs, which is advantageous to recall. In this case, methods like DEGseq stand out as they can maintain recall while not sacrificing precision since it will tend to call more genes as DEGs on average compared to its counterparts. DEGseq also operates invariantly at FDRs of 5–20%, making it highly suitable for precision medicine since an FDR of 5% is a default standard in clinical decision-making. In the overly conservative scenario with smaller number of DEGs in the gold standard, a more selective approach will perform better, highlighted in the precision parameter and illustrating the trade-offs available across all the tested techniques. An ensemble provides the analyst a robust trade-off alternative as it can build upon the strengths of all methods, and not suffer the issue of “performing well” in one dataset but not in another. In each panel, methods are ordered according to performance.

**Figure 5.**
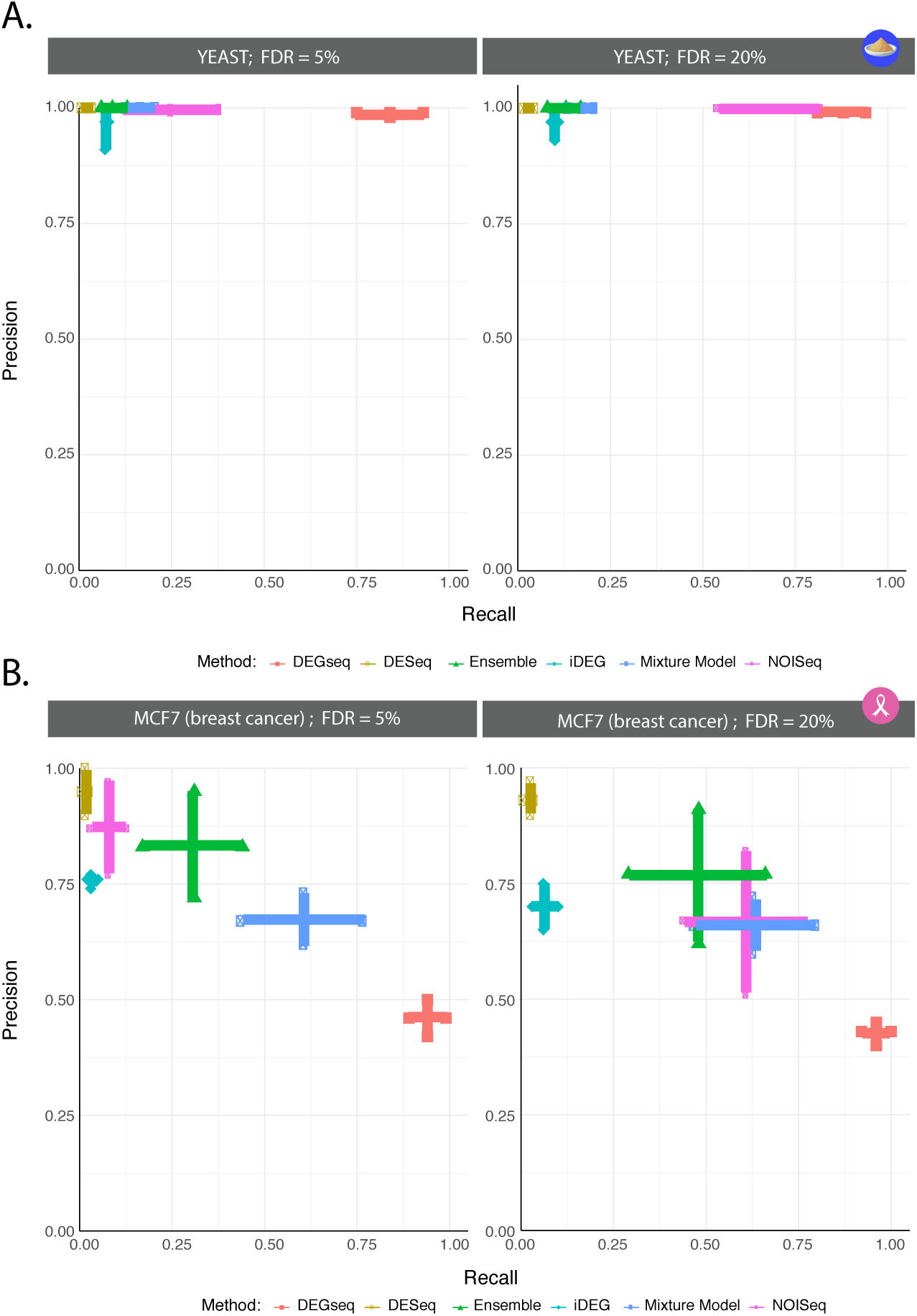
The optimal Precision-Recall summary plots in Yeast and MCF7 breast cancer cell lines were constructed by summarizing precision-recall confidence regions over every ss-DEG evaluation studies and indicating the broadest possible precision-recall combinations. The summarized curves show a spectrum of operating characteristics and performance across techniques, indicating the need for an ensemble-like approach and substantial improvements in ss-DEG. The MCF7 case study produced reference standards that predicted between 15% of the genes in the genome as DEGs, while the Yeast case study produced reference standards that predicted between 55% and 70% of the genes in the genome as DEG. The more clinically relevant range of DEGs from the MCF7 reference standard construction introduces a very distinct detection problem where methods like DEGseq result in a large number of False Positive as shown in the precision-recall summary plots. It achieves high recall at the expense of low-precision. Conversely, conservative techniques like DESeq obtain a very high precision on a small number of calls. The results show that this is indeed a challenging detection task across the board, and that various techniques operate differently, providing an analyst with a wide-range of operating characteristics. In the Yeast dataset, all methods achieve a high precision, with varying levels of recall, however given that the majority of genes are labeled DEGs, this favors methods with high number of calls. The MCF7 breast cancer cell line analysis shows that in lower-signal settings, an ensemble approach optimizes the precision-recall trade-off. Since certain methods can perform well in one scenario and underperform in others, we recommend a contextual use or an ensemble-like approach where the strengths of these tools can be combined into a single, robust predictor. Here, precision and recall of each instance of ss-DEGs are respectively calculated on the union and the intersection of reference standards (**Table 2**). Recall and precision are reported as one standard deviation in each direction creating a cross. Of note, at FDR<20%, DESeq2 produces no predictions and is thus not shown and considered inappropriate for single-subject DEG analyses.

## Discussion

Our analyses clearly demonstrated the intricacies of working with biologically complex transcriptomic data in the absence of ground truth. As shown by **Figures 4** and **5**, NOISeq-sim outperformed other tested single-subject techniques in terms of precision across both case studies, and was capable of scoring well across a range of cohort-derived reference standards. In contrast, single-subject implementations of DESeq and DESeq2 were highly conservative, with ss-DESeq2 predicting zero DEGs, regardless of dataset, it doesn’t seem to perform without replicate in real datasets. Our ensemble model was able to obtain consistent high overall accuracies which suggests that a combination of parameter and distribution assumptions can overcome some of the limitations and biases inherent to any one model and approach a more accurate consensus standard (Fig 4A).

From this study, it also appears that all ss-DEG methods are sensitive to the percentage of DEGs present in the reference set. Given this, the degree of perturbation, or, range in number of DEGs expected in a pair of samples can guide the method selection. The Yeast dataset was utilized because of its large number of replicates to construct independent test and validation sets, however, the range of DEGs observed as a consequence of deleting a component of the transcriptional machinery is clearly higher than expected between most paired clinical samples. On the other hand, the MCF7 dataset was limited in term of samples while providing some insight in DEG ranges of 15%-30%. We had no datasets to evaluate conditions with DEGs<15%. As simulations and synthetic data can investigate a ranges of accuracies against a true gold standard, though they can be prone to other biases and limitations. Li et al. (14) have implemented a comprehensive simulation of ss-DEG methods across 8000 tests in a companion study, using a range of DEG proportions from 5–40%, assuming distinct distributions (Poisson or negative binomial), and modeling a variable mean to variance relationship observed from real datasets as recommended by McCarthy et al. (32). The results from those simulations broadly agree with the results obtained in this study, identifying the same precision and recall rankings between NOISeq-sim, edgeR, DESeq, and DEGSeq. In contrast, however, simulation studies generally yielded higher recall estimates, which suggests the observed residual cross-replicate heterogeneity comprised of non-genomic and stochastic variation of real biologic datasets can substantially limit performance of the DEG methods applied to two conditions without replicates. Due to this, we suggest that these methods’ performance should be seen as a range or spectrum, rather than definite.

Based on the results shown in **Figures 4** and **5**, we report in **Table 4** recommendations for the use of ss-DEGs in two conditions without replicates. Of note, when comparing our results to the performance metrics published alongside the Yeast and MCF7 data in the original publications by Schurch(18) and Liu(19), we found that the performance was lower across our studies. One reason for this is that those authors calculated the accuracy of their r-DEG methods in the presence of replicates using anticonservative conditions: each method was compared to itself using the total number of replicates, while sub studies utilized a random sample within those utilized for the reference standard. Here, our accuracies were more conservatively calculated in two ways: (i) we constructed each gold standard by using distinct samples for predictions without predicates from the reference standard construction, and (ii) the accuracy scores of a method predicting DEGs without replicates were tested against reference standards built by distinct methods in replicates.

**Table 4.**
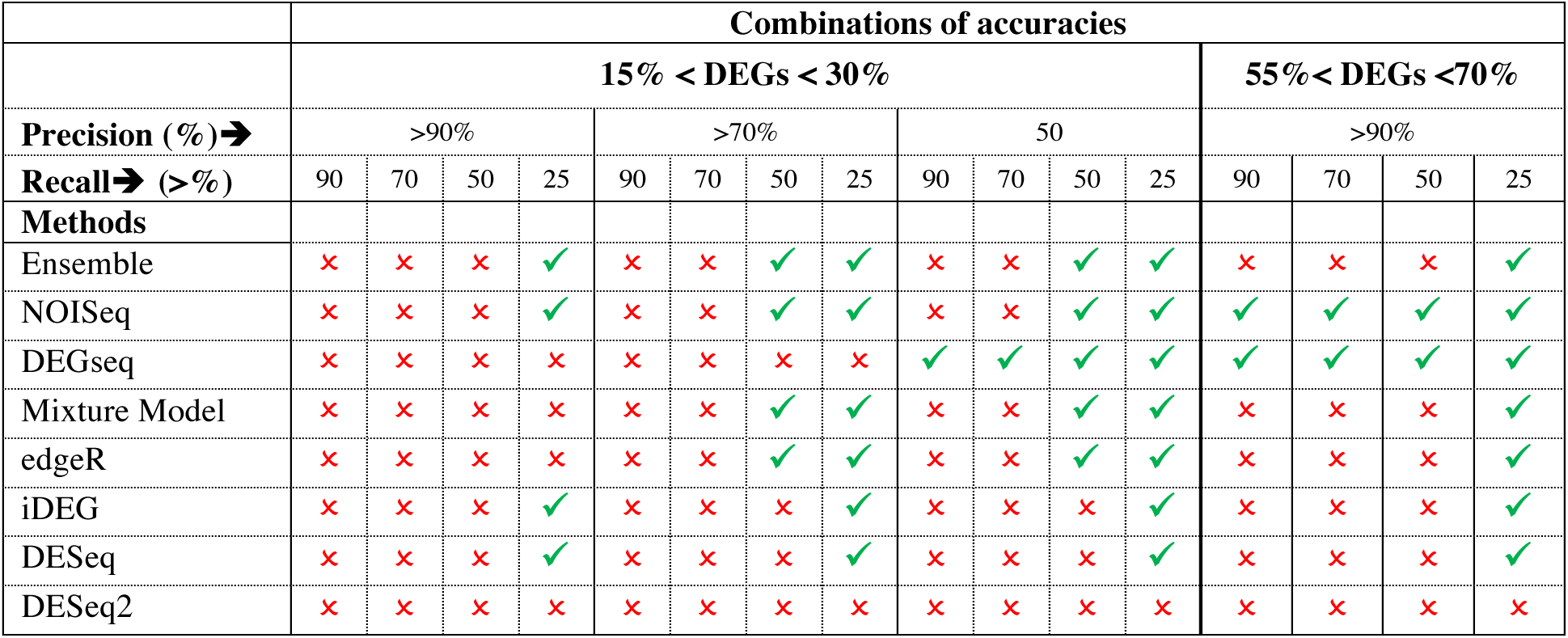
Recommendations on ss-DEG methods applied to single-subject studies of two-sample conditions without replicates. *✓* = recommended; ✗ = not recommended

**Figure 4A** shows how DEGseq performs similarly to NOISeq and the ensemble method maintains precision at FDR 5% and properly detects nearly 75% of the DEGs. One could argue this combination would potentially make DEGseq the ideal tool for this dataset; however, in the MCF7 case study, DEGseq could not replicate its performance (**Figure 5b**). This highlights the risks of relying on a single technique across distinct DEG proportions. Furthermore, under multiple biological replicates, techniques like DESeq, DESeq2, and edgeR are the staples of RNA-Seq data analysis and are often the authors’ default method choices and recommendations for building reference standards. However, as seen in **Figures 4** and **5**, DESeq and DESeq2 performed conservatively in the two datasets, showing that their reliability does not necessarily extend to N-of-1 and no-replicate conditions. This study provided a promising first comparison of how ss-and r-DEGs RNA-Seq techniques fare in absence of replicates, and further datasets and analyses will better confirm or clarify these findings.

Another promising result from this study came from the implementation of an ensemble approach which has the potential to provide a robust, stabilizing aggregation that can perform as well (if not better) as individual techniques without the pitfalls of inconsistently performing across datasets. Therefore, in absence of knowing the truth or having a reliable gold standard, we would recommend an ensemble-like approach. The Machine Learning community has shown ensemble learners and classifiers to perform well in a variety of settings. Since it has been shown that diversity helps ensemble classifiers (33), the ensemble was constructed by averaging the predictions from DEGseq, NOISeq, Mixture Models, and edgeR. Since these four methods each assume different stochastic data models (Binomial, Non-parametric, Mixture Model, and Negative Binomial, respectively), they essentially capture the distinct flavors of the available techniques and provide the ensemble with the consensus of computationally derived ‘expert opinions’.

In addition, ranges of DEGs< 15% merit to be explored as they are likely clinically relevant in response to therapy. The addition of more datasets with additional response ranges would further improve our understanding of the accuracy of ss-DEG methods, especially when these datasets are previously validated, as was the case of the MCF7 and Yeast datasets. Furthermore, improvement of ss-DEGs methods is required, particularly for performing with higher recall when DEGs are low. Further studies are also needed to describe the effectiveness of better performing methods, such as NOISeq (specifically NOISeq-sim), in the absence of replicates across different contexts.

In the past, comparing a transcriptome to heterogenic samples from other subjects has been proposed. However, this strategy brings up a number of confounding factors: distinct genetics, distinct environmental factors, etc. Here, we proposed using one’s own samples as controls. Adding biological replicates increases accuracy and is recommended where possible, but may not be feasible in certain clinical settings and can be cost prohibitive. In the absence of replicates, focusing on identifying those DEGs within differentially expressed pathways may further improve the accuracy rates and merit validation in future studies.

## Conclusions

This study demonstrates that determining differentially expressed genes (DEGs) between two samples of one subject can be obtained with high precision and limited recall (<30%) when the true number of DEGs ranges from 15%-30%, while a few methods can provide reliable results under conditions where the proportion of DEGs exceed 50% of the genome. No single ss-DEG method obtained both high precision and recall in the evaluations using these biological datasets, though some obtained a reasonably robust F1-score.

As RNA-Seq technologies expand the scope and problem space into single-subject and N-of-1 settings and data analysis tools keep growing, more time and research need to focus on a greater understanding of which analysis tools are better suited for clinical samples and individual inference. At the moment, the limited access to a sufficient quantity of clinically relevant tissues typically prohibits replicates. Thus, analysis methods that require replicates to determine DEGs must be adapted or replaced in order to advance the utility of transcriptome profiling in precision medicine. Through the work in this study, we show that ongoing improvements in single-subject methods are required for these to work robustly and accurately in absence of replicates. We have also shown that the biological and data characteristics of RNA-seq comparison are also critical factors for method performance, as the relative strengths and limitations of each method differed markedly depending on the proportion of DEGs regulated by the bioassay. RNA-Seq datasets cannot and should not be treated as a monolith with a universal analysis standard applied, ignoring specific properties and subject matter expectations.

We also show that it remains difficult to generate consensus reference standards from different RNA-seq analysis tools as the intersection of all methods agreed on less than 50% of called DEGs, even while implementing these tools under their recommended conditions with replicate samples.

In order to improve the accuracy, we propose that future methods consider the injection of knowledge from curated gene set (e.g. Gene Ontology) and network science (e.g., unbiased gene set obtained from co-expression networks) to pool the signal of altered genes belonging to functional units as a way to increase signal accuracy and reliability. While the reductionism of identifying directly DEGs is appealing, previous systems genomics work showing stronger signals at differentially expressed pathways in single-subject studies suggest combining the two approaches would substantially increase accuracies.

## List of Abbreviations

- ss = Single-Subject
- r = cohort of isogenic replicates-based
- DEG= Differentially Expressed Genes
- FDR = False Discovery Rate (specifically, BY)
- WT = Wild Type
- AUC = Area Under the Curve
- PR = Precision-Recall
- SD = Standard Deviation

## Authors’ contributions

SRZ conducted all the analyses in R; YAL and JB contributed to the interpretation of the study; CK, YAL, and SRZ contributed to the figures and tables; SRZ, CK, JB, and YAL contributed to the writing of the manuscript; all authors read and approved the final manuscript.

## Acknowledgements

We thank Dr. Qike Li for providing access of his iDEG program, PhD Thesis and manuscript under review. We thank Mr. Dillon Aberasturi for conducting preliminary studies.

**Supplement Table S1.** Methods comparing gene sets and pathways between two transcriptomes that have been reproducibly demonstrated as feasible in single subjects (3).

| Method | Transcript P-Value Call | Pathway P-value | Validation of application to single subject |  |  |
| --- | --- | --- | --- | --- | --- |
|  |  |  | *Internal* |  | *External* |
|  |  |  | Simulation | Biological Replicates or Gold Standard | Translation to diagnosis, prognosis & treatment |
| FAIME (34) | ✗ | ✗ | ✓ | ✓ | ✓ |
| N-of-1 Pathway Wilcoxon (6) | ✗ | ✓ | ✓ | ✓ | ✓ |
| N-of-1 Pathways MixEnrich (5) | ✓ | ✓ | ✓ | ✓ | ✓ |
| N-of-1 Pathways kMen (25) | ✗ | ✓ | ✓ | ✓ | ✓ |
| N-of-1 Pathways MD (7) | ✗ | ✓ | ✓ | ✓ | ✓ |
| N-of-1 Pathways Wilcoxon (6) | ✗ | ✓ | ✓ | ✓ | ✓ |

## Footnotes

i Data downloaded from on August 27^th^, 2018. https://github.com/bartongroup/profDGE48

ii NOISeq-Bio was used to construct the reference standard, while NOISeq-sim was used in the single-subject prediction sets.

iii Partial validation conducted using qPCR with 400 genes with ~ 80% DEGs.

iv Mixture Model provides a posterior probability rather than a p-value, when FDR<5% is indicated in the manuscript, it translates as a posterior probability >95% for the mixture models.

## References

1. Buguliskis JS. Could RNA-Seq Become the Workhorse of Precision Medicine? Genetic Engineering & Biotechnology News. 2015 March 01, 2015.

2. Holik AZ, Law CW, Liu R, Wang Z, Wang W, Ahn J, et al. RNA-seq mixology: designing realistic control experiments to compare protocols and analysis methods. Nucleic Acids Res. 2017;45(5):e30.

3. Vitali F, Li Q, Schissler AG, Berghout J, Kenost C, Lussier YA. Developing a ‘personalome’ for precision medicine: emerging methods that compute interpretable effect sizes from single-subject transcriptomes. Brief Bioinform. 2017.

4. Ozturk K, Dow M, Carlin DE, Bejar R, Carter H. The Emerging Potential for Network Analysis to Inform Precision Cancer Medicine. J Mol Biol. 2018;430(18 Pt A):2875–99.

5. Li Q, Schissler AG, Gardeux V, Achour I, Kenost C, Berghout J, et al. N-of-1-pathways MixEnrich: advancing precision medicine via single-subject analysis in discovering dynamic changes of transcriptomes. BMC Medical Genomics. 2017;10(1):27.

6. Gardeux V, Achour I, Li J, Maienschein-Cline M, Li H, Pesce L, et al. ‘N-of-1-pathways’ unveils personal deregulated mechanisms from a single pair of RNA-Seq samples: towards precision medicine. Journal of the American Medical Informatics Association. 2014;21(6): 1015–25.

7. Schissler AG, Gardeux V, Li Q, Achour I, Li H, Piegorsch WW, et al. Dynamic changes of RNA-sequencing expression for precision medicine: N-of-1-pathways Mahalanobis distance within pathways of single subjects predicts breast cancer survival. Bioinformatics. 2015;31(12):i293–i302.

8. Schissler AG, Li Q, Chen JL, Kenost C, Achour I, Billheimer DD, et al. Analysis of aggregated cell-cell statistical distances within pathways unveils therapeutic-resistance mechanisms in circulating tumor cells. Bioinformatics. 2016;32(12):i80–i9.

9. Schissler AG, Piegorsch WW, Lussier YA. Testing for differentially expressed genetic pathways with single-subject N-of-1 data in the presence of inter-gene correlation. Stat Methods Med Res. 2017: 962280217712271.

10. Li Q, Schissler AG, Gardeux V, Berghout J, Achour I, Kenost C, et al. kMEn: Analyzing noisy and bidirectional transcriptional pathway responses in single subjects. J Biomed Inform. 2017;66:32–41.

11. Gardeux V, Arslan AD, Achour I, Ho TT, Beck WT, Lussier YA. Concordance of deregulated mechanisms unveiled in underpowered experiments: PTBP1 knockdown case study. BMC Med Genomics. 2014;7 Suppl 1:S1.

12. Gardeux V, Bosco A, Li J, Halonen MJ, Jackson D, Martinez FD, et al. Towards a PBMC “virogram assay” for precision medicine: Concordance between ex vivo and in vivo viral infection transcriptomes. Journal of biomedical informatics. 2015;55:94–103.

13. Gardeux V, Berghout J, Achour I, Schissler AG, Li Q, Kenost C, et al. A genome-by-environment interaction classifier for precision medicine: personal transcriptome response to rhinovirus identifies children prone to asthma exacerbations. Journal of the American Medical Informatics Association. 2017:ocx069.

14. Li Q, Zaim SR, Aberasturi D, Berghout J, Li H, Kenost C, et al. iDEG: a single-subject method utilizing local estimates of dispersion to impute differential expression between two transcriptomes. *bioRxiv*. 2018. https://doi.org/10.1101/405332

15. Tarazona S, García F, Ferrer A, Dopazo J, Conesa A. NOIseq: a RNA-seq differential expression method robust for sequencing depth biases. EMBnet journal. 2011;17(B):18–9.

16. Li X, Brock GN, Rouchka EC, Cooper NG, Wu D, O’Toole TE, et al. A comparison of per sample global scaling and per gene normalization methods for differential expression analysis of RNA-seq data. PloS one. 2017;12(5):e0176185.

17. Team RC. R: A language and environment for statistical computing. 2013.

18. Schurch NJ, Schofield P, Gierliński M, Cole C, Sherstnev A, Singh V, et al. How many biological replicates are needed in an RNA-seq experiment and which differential expression tool should you use? RNA. 2016;22(6):839–51.

19. Liu Y, Zhou J, White KP. RNA-seq differential expression studies: more sequence or more replication? Bioinformatics. 2014;30(3):301–4.

20. Edgar R, Domrachev M, Lash AE. Gene Expression Omnibus: NCBI gene expression and hybridization array data repository. Nucleic acids research. 2002;30(1):207–10.

21. Robinson MD, McCarthy DJ, Smyth GK. edgeR: a Bioconductor Package for Differential Expression Analysis of Digital Gene Expression Data. Bioinformatics. 2009;26(1): 139–40.

22. Anders S, Huber W. Differential expression of RNA-Seq data at the gene level-the DESeq package. Heidelberg, Germany: European Molecular Biology Laboratory (EMBL). 2012.

23. Love MI, Huber W, Anders S. Moderated estimation of fold change and dispersion for RNA-seq data with DESeq2. Genome biology. 2014; 15(12):550.

24. Wang L, Feng Z, Wang X, Wang X, Zhang X. Degseq: an R Package for Identifying Differentially Expressed Genes From Rna-Seq Data. Bioinformatics. 2009;26(1): 136–8.

25. Li Q, Schissler AG, Gardeux V, Berghout J, Achour I, Kenost C, et al. kMEn: Analyzing noisy and bidirectional transcriptional pathway responses in single subjects. Journal of biomedical informatics. 2017;66:32–41.

26. Benjamini Y, Yekutieli D. The control of the false discovery rate in multiple testing under dependency. Ann Stat. 2001;29(4):1165–88.

27. Breiman L. Random Forests. Machine Learning. 2001;45(1):5–32.

28. Benjamini Y, Yekutieli D. The Control of the False Discovery Rate in Multiple Testing under Dependency. The Annals of Statistics. 2001;29(4):1165–88.

29. Saito T, Rehmsmeier M. Precrec: fast and accurate precision-recall and ROC curve calculations in R. Bioinformatics. 2017;33(1): 145–7.

30. Wickham H, Chang W. ggplot2: An implementation of the Grammar of Graphics. R package version 07, URL: http://CRANR-projectorg/package=ggplot2. 2008.

31. Bullard JH, Purdom E, Hansen KD, Dudoit S. Evaluation of statistical methods for normalization and differential expression in mRNA-Seq experiments. BMC Bioinformatics. 2010;11:94.

32. McCarthy DJ, Chen Y, Smyth GK. Differential expression analysis of multifactor RNA-Seq experiments with respect to biological variation. Nucleic Acids Res. 2012;40.

33. Kuncheva LI, Whitaker CJ. Measures of diversity in classifier ensembles and their relationship with the ensemble accuracy. Machine learning. 2003;51(2):181–207.

34. Yang X, Regan K, Huang Y, Zhang Q, Li J, Seiwert TY, et al. Single sample expression-anchored mechanisms predict survival in head and neck cancer. PLoS computational biology. 2012;8(1):e1002350.

